# Intranasal gene therapy to prevent infection by SARS-CoV-2 variants

**DOI:** 10.1101/2021.04.09.439149

**Authors:** Joshua J. Sims, Jenny A. Greig, Kristofer T. Michalson, Sharon Lian, R. Alexander Martino, Rosemary Meggersee, Kevin B. Turner, Kalyani Nambiar, Cecilia Dyer, Christian Hinderer, Makoto Horiuchi, Hanying Yan, Xin Huang, Shu-Jen Chen, James M. Wilson

**Affiliations:** Gene Therapy Program, Department of Medicine, Perelman School of Medicine, University of Pennsylvania, Philadelphia, PA 19104, USA

## Abstract

SARS-CoV-2 variants have emerged with enhanced pathogenicity and transmissibility, and escape from pre-existing immunity, suggesting first-generation vaccines and monoclonal antibodies may now be less effective. This manuscript demonstrates an approach for preventing clinical sequelae and the spread of SARS-CoV-2 variants. First, we affinity-matured an angiotensin-converting enzyme 2 (ACE2) decoy protein, achieving 1000-fold binding improvements that extend across a wide range of SARS-CoV-2 variants and distantly related, ACE2-dependent coronaviruses. Next, we demonstrated the expression of this decoy in proximal airway when delivered via intranasal administration of an AAV vector. This intervention significantly diminished clinical and pathologic consequences of SARS-CoV-2 challenge in a mouse model and achieved therapeutic levels of decoy expression at the surface of proximal airways when delivered intranasally to nonhuman primates. Importantly, this long-lasting, passive protection approach is applicable in vulnerable populations such as the elderly and immune-compromised that do not respond well to traditional vaccination. This approach could be useful in combating COVID-19 surges caused by SARS-CoV-2 variants and should be considered as a countermeasure to future pandemics caused by pre-emergent members, ACE2-dependent CoVs that are poised for zoonosis.

**Author summary:** SARS-CoV-2 variants have emerged with enhanced pathogenicity and transmissibility, and escape from pre-existing immunity, suggesting first-generation vaccines and monoclonal antibodies may now be less effective. This manuscript demonstrates an approach for preventing clinical sequelae and the spread of SARS-CoV-2 variants. First, we affinity-matured an angiotensin-converting enzyme 2 (ACE2) decoy protein, achieving 1000-fold binding improvements that extend across a wide range of SARS-CoV-2 variants and distantly related, ACE2-dependent coronaviruses. Next, we demonstrated the expression of this decoy in proximal airway when delivered via intranasal administration of an AAV vector. This intervention significantly diminished clinical and pathologic consequences of SARS-CoV-2 challenge in a mouse model and achieved therapeutic levels of decoy expression at the surface of proximal airways when delivered intranasally to nonhuman primates. Importantly, this long-lasting, passive protection approach is applicable in vulnerable populations such as the elderly and immune-compromised that do not respond well to traditional vaccination. This approach could be useful in combating COVID-19 surges caused by SARS-CoV-2 variants and should be considered as a countermeasure to future pandemics caused by pre-emergent members, ACE2-dependent CoVs that are poised for zoonosis.

## Introduction

Developing a soluble form of angiotensin-converting enzyme 2 (ACE2)—referred to as a decoy—is considered a protein therapeutic in the treatment of COVID-19 patients (1, 2). We isolated an ACE2 decoy that broadly neutralizes SARS-CoV-2 variants and demonstrate its potential for preventing COVID-19 when expressed from an adeno-associated virus (AAV) following intranasal (IN) delivery. We have reported previously on the effectiveness of IN AAV to express antibodies that broadly neutralize pandemic strains of influenza (3-6).

## Results

### ACE2 decoy affinity maturation enhances neutralization of SARS-CoV-2 100-fold

We initially constructed a decoy receptor by fusing a human ACE2 fragment to the human IgG4 Fc domain. We cloned this first-generation decoy into AAV and delivered it as a nasal spray into nonhuman primates (NHPs). Although we detected decoy expression in nasal lavage fluid (NLF), the decoy was not produced at levels sufficient to overcome the low neutralizing potency of this protein (Figure S1). We therefore set out to affinity-mature the ACE2 protein sequence.

We generated diverse (>10^8^ transformants) ACE2 variant libraries in a yeast-display format(7) using error-prone polymerase chain reaction (PCR; Figure 1A and Figure S2). We screened the primary libraries in rounds of fluorescence-activated cell sorting (FACS; Figure 1B). We selected populations with better binding to SARS-CoV-2 receptor binding domain (RBD) and tracked library convergence with deep sequencing (Figure 1C and 1D). Frequently observed mutations from our primary library sorts overlap partially with mutations reported by others(2, 8), including substitutions at T27 and N90 glycan disruption (Figure S2). Validated clones from the sorted primary libraries (Figure S3) seeded a secondary library formed by mutagenic recombination(9, 10), which we screened using stringent off-rate sorting(11) (Figure 1C).

**Fig. 1.**
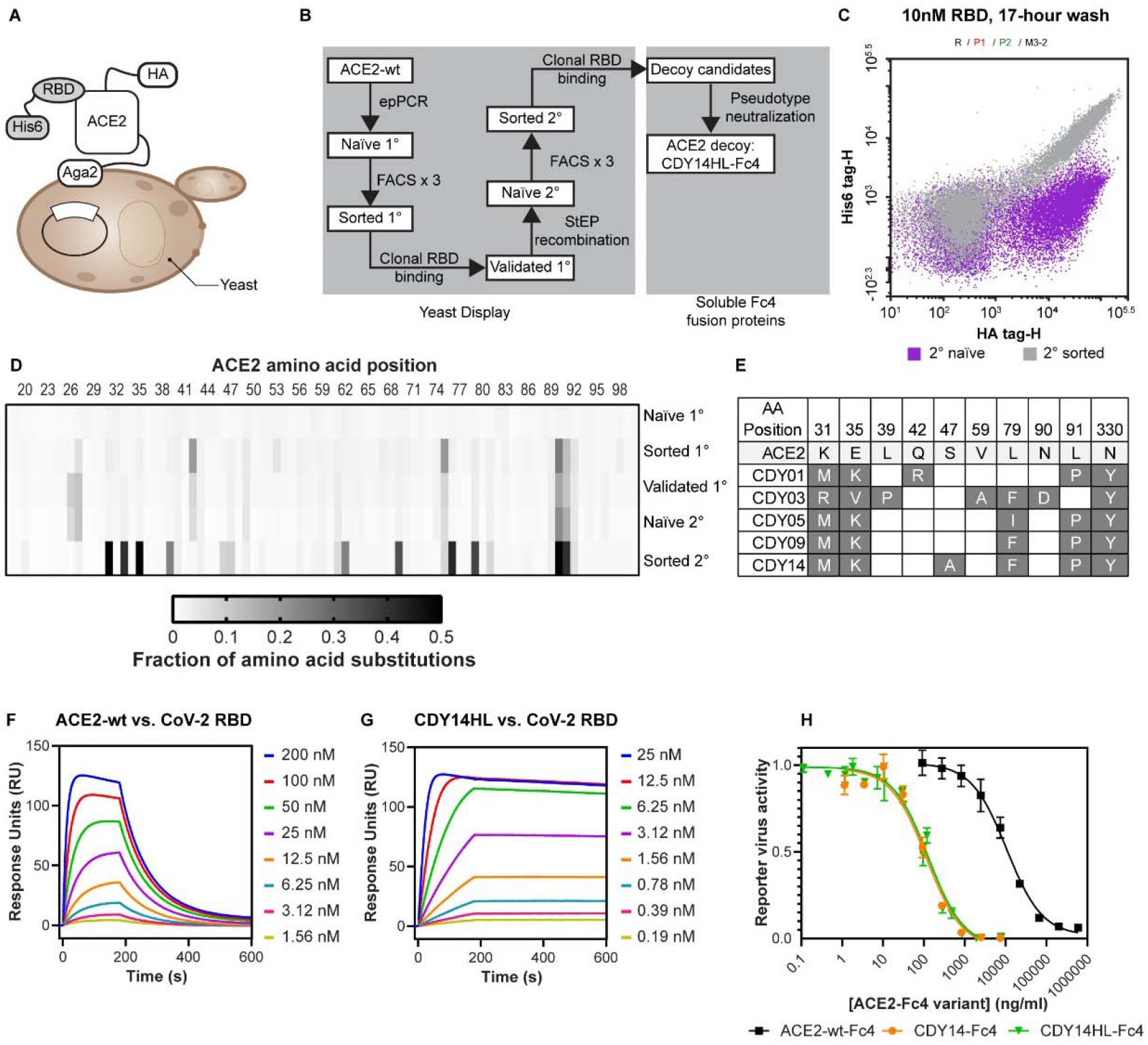
ACE2 Decoy Receptor Engineering. **A**. Yeast display (YD) system **B**. Decoy affinity maturation and candidate selection process. **C**. Flow cytometry analysis of the naïve (purple) and sorted populations (gray) from the secondary YD library. **D**. NGS analysis of plasmid populations recovered from rounds of YD. **E**. Mutations accumulated in the top five decoy-Fc4 candidates. **F and G**. SPR binding analysis for CoV-2 RBD injected over surface-immobilized ACE2-wt (F) or CDY14HL (G). **H**. Wuhan CoV-2-Pseudotyped lentiviral reporter neutralization assay of ACE2-wt-Fc4, CDY14-Fc4, and CDY14HL-Fc4. Data for at least three independent measurements are presented as average ± standard deviation.

We expressed ACE2 variants from several stages of the yeast-display screening as soluble IgG4 Fc fusions, evaluated expression titers, and predicted IC_50_ for SARS-CoV-2 neutralization using reporter virus (Figure S3). The most potent neutralizing variants converged upon similar substitutions at five positions: 31, 35, 79, 330, and N90 glycan disruption (Figure 1E). After further characterization, we selected CDY14-Fc4 as the most improved ACE2 decoy variant. To avoid off-target effects *in vivo*, we ablated ACE2 enzyme activity by introducing H345L(12) at no cost to potency. By surface plasmon resonance (SPR) the active site-null CDY14HL-Fc4 bound SARS-CoV-2 RBD with 1,000-fold improved affinity (29 nM for wtACE2 vs. 31 pM for CDY14HL-Fc4; see Figure 1, panels F and G). CDY14HL-Fc4 neutralized Wuhan-Hu-1 SARS-CoV-2 reporter nearly 100-fold better than the un-engineered ACE2 decoy (IC_50_ 127 ng/ml for CDY14HL-Fc4 vs. 11 µg/ml for ACE2-wt-Fc4; see Figure 1H).

### ACE2 decoy is effective against SARS-CoV-2 variants and SARS-CoV-1

Escape mutations at the immunodominant ACE2 binding site of the RBD is of major concern for emerging SARS-CoV-2 variants(13). RBDs from more distant ACE2-dependent CoVs also differ substantially from the original SARS-CoV-2 at the ACE2 interface (Figure 2A). Unlike antibodies, decoy inhibitors may achieve broad neutralization and escape mutant resistance; changes that reduce decoy binding would also decrease ACE2 receptor binding, thus reducing viral fitness. To assess this potential, we measured ACE2-Fc4 (a surrogate for the native receptor) and CDY14HL-Fc4 (the therapeutic decoy) affinities across a diverse panel of CoV RBDs using SPR. We chose RBDs from strains under positive selection in the course of the 2020 pandemic (e.g., 439K (14), B.1.1.7 and B1.351, first isolated in the EU, UK, and the Republic of South Africa, respectively(15)), and mink-adapted isolates(16). Several emerging SARS-CoV-2 variants with improved affinity for ACE2-Fc4 (B1.1.7, 453F, and 501T) also bind CDY14HL-Fc4 more tightly (Figure 2B and Figure S4), while B.1.351, 439K, and 439K/417V only modestly alter binding to ACE2-Fc4 or CDY14HL-Fc4. While 486L reduces affinity for CDY14HL-Fc4, it does so for ACE2-Fc4 proportionally. Remarkably, decoy and receptor RBD affinities are tightly coupled even for the distantly related SARS-pandemic CoV-1 and the pre-emergent bat WIV1-CoV(17).

**Fig. 2.**
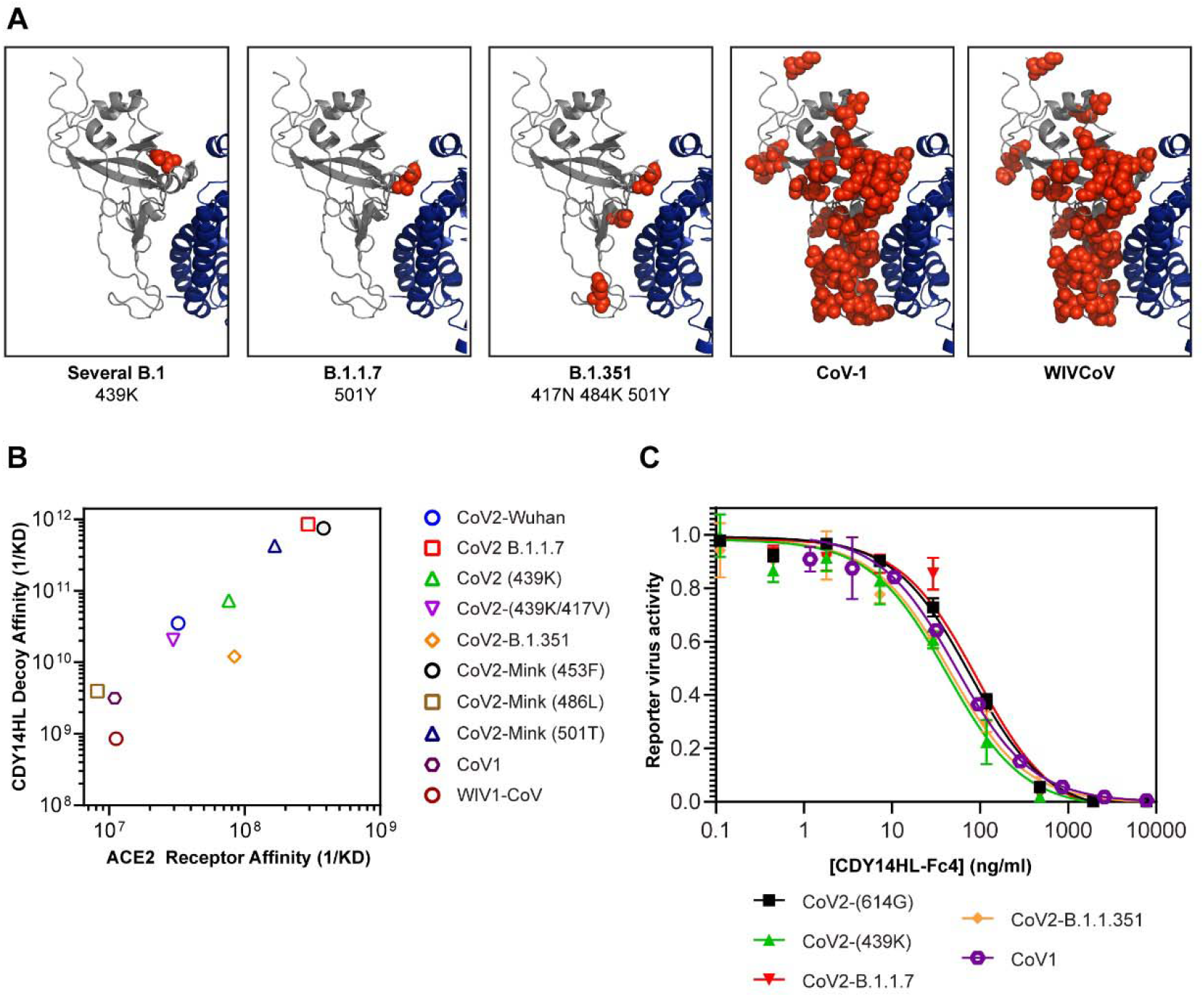
ACE2 Decoy Binding and Neutralization Across Diverse CoVs. **A**Structural models (7DF4.pdb (40)) of Wuhan CoV-2 RBD (gray) bound to human ACE2 (blue) RBD residues that distinguish each variant (listed below each panel) from the Wuhan CoV-2 RBD are shown in red spheres. **B**. SPR measurements: ACE2-wt-Fc4 or CDY14HL-Fc4 binding to various purified recombinant RBD proteins. **C**. We used reporter lentiviruses pseudotyped with CoV spike proteins from five isolates to determine the neutralizing potencies of CDY14HL-Fc4. Reporter virus activity data are presented as mean ± standard deviation for at least three replicate titrations.

Next, we compared decoy neutralization across diverse SARS-CoVs using pseudotyped lentivirus reporters. The SARS-CoV-2 variant reporter viruses in the 614G (18, 19) background (439K, B1.1.7, B.1.351) are neutralized near or below the IC_50_ of the 614G reporter (IC_50_ values: 77 ng/ml for 614G, 42 ng/ml for 439K, 93 ng/ml for B.1.1.7, and 45 for B.1.351; Figure 2C). Remarkably, selecting for increased binding to the SARS-CoV-2 RBD resulted in very potent neutralization of the phylogenetically distinct SARS-CoV-1 reporter virus (IC_50_ = 53 ng/ml). Taken together with the binding survey, these data indicate that structural features of the ACE2 interface have been retained through the stages of directed evolution. Moreover, the data predict that CDY14HL-Fc4 could protect against current, emerging, and future pandemic ACE2-dependent CoVs.

### ACE2 decoy diminishes SARS-CoV-2 sequelae in transgenic mice

We considered SARS-CoV-2 challenge studies in hamsters, macaques, and the hACE2 transgenic (TG) mice to evaluate the *in vivo* efficacy of an AAV vector expressing CDY14HL-Fc4. In all models, achieving evidence of viral replication *in vivo* requires virus doses that far exceed those required for human transmission. Furthermore, the clinical and pathologic sequalae of SARS-CoV-2 exposure is attenuated in these species compared to severely affected humans. The most significant limitation, however, is that all the challenge models require direct pathogen delivery to the lung in order to demonstrate pathology, which does not simulate the mechanism of the AAV decoy product, which focuses on localized expression in the proximal airway following intranasal delivery to reduce SARS-CoV-2 infection and its consequences. We therefore selected the hACE2 TG mouse model for three reasons: 1) we can characterize disease by measuring viral loads, clinical sequalae, and histopathology; 2) we can use an IN route of administration as we would in humans, realizing this deposits vector in the proximal and distal airways of the mouse, while IN delivery in humans is restricted to the proximal airway; and 3) we can leverage the extensive experience of murine models in de-risking human studies of AAV gene transfer.

We conducted pilot studies in wild-type mice to determine which decoy protein (CDY14-Fc4 vs. CDY14HL-Fc4) and capsid (clade F AAVhu68 vs. clade A AAVrh91) maximized expression following *in vivo* gene delivery. We administered 10^11^ GC of vector and recovered broncho-alveolar lavage samples (BAL) 7 days later to evaluate ACE2 decoy protein expression and activity (Figure 3A-C). Based on mass spectrometry (MS) protein measurements, the AAVhu68 capsid was more efficient than the AAVrh91 capsid in transducing mouse lung. The HL mutation modestly reduced expression (p<0.007). Importantly, we found a direct correlation between decoy expression levels and the ability to bind to SARS-CoV-2 spike protein and neutralize a SARS-CoV-2 pseudotype, demonstrating function of the decoy expressed from airway tissues (Figure 3A-C). We selected CDY14HL-Fc4 as the clinical candidate transgene and the AAVhu68 capsid for the mouse challenge studies (Figure 3D).

**Figure 3.**
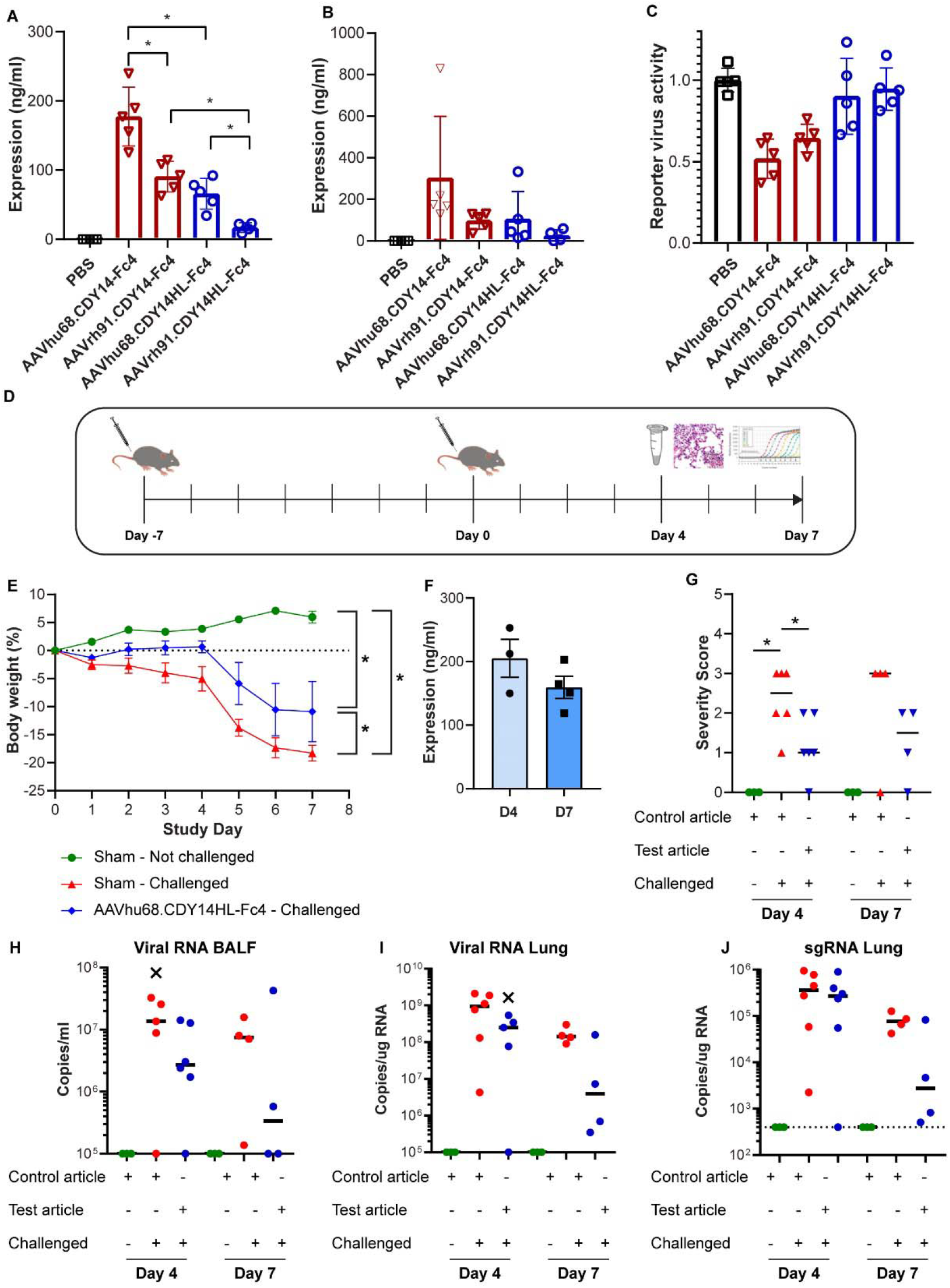
Protection in the human ACE2 transgenic mouse model. BAL from vector-treated animals analyzed for: **A**. decoy protein by MS; **B**. SARS-CoV-2 spike ELISA; **C**. neutralization of SARS-CoV-2 pseudotyped lentivirus. **D**. Challenge study design. **E**. Weight loss in the animals that were sustained for 7 days; one animal in the vehicle and vector treated groups required euthanasia. **F**. MS assay of expression in ASF (corrected for BAL dilution). **G**. Pulmonary inflammation histopathology scores of tissues harvested at days 4 and 7. **H**. Viral RNA in BAL. **I**. Viral RNA in lung. **J**. Sub-genomic RNA in lung. Outliers are indicated with X.

We IN delivered AAVhu68-CDY14HL-Fc4 or vehicle to hACE2-TG mice. Seven days later, animals were challenged with SARS-CoV-2 (280 pfu), followed clinically (observation and daily weights), and necropsied on days 4 and 7 after challenge for tissue and BAL analysis (Figure 3D). Expression of CDY14HL-Fc4 in BAL normalized for dilution was in the range of the IC_50_ measured *in vitro* and in the pilot studies (Figure 3F). Sham-treated SARS-CoV-2 challenged animals demonstrated statistically significant weight loss as has been described by others(20-22). We observed significantly less weight loss amongst vector-treated animals (which we followed for 7 days) compared to untreated animals (observed on days 4 and 7; p<0.05, linear mixed effect modeling). The vector-treated animals also significantly differed from the untreated, unchallenged animals (Figure 3E). Interestingly, the clinical outcome of the treatment was better among females than males, although we noted significant variations within the treated group (Figure S5).

Histopathology of the lungs from vehicle treated animals challenged with SARS-CoV-2 revealed findings similar to that previously described in this model(22). As expected, tissues from animals not challenged with SARS-CoV-2 demonstrated no histopathology. Samples from days 4 and 7 showed reduced lung pathology in AAVhu68.CDY14HL-Fc4 treated animals vs. the vehicle-treated animals; the day-4 samples achieved statistical significance (p<0.05; Wilcoxon Rank Sum Test). (Figure 3G). Compared to vehicle-treated animals, viral RNA in BAL and lung homogenate was diminished at day 4 and 7 in AAVhu68.CDY14HL-Fc4 treated animals (Figure 3H and 3I). The greatest reductions were at day 7 for both BAL (26-fold) and lung tissue (35-fold). Impact of the AAVhu68.CDY14HL-Fc4 on SARS-CoV-2 replication, as determined by median sgRNA levels, was greatest at day 7 (27-fold reduction, Figure 3J). Although there was substantial inter-animal variation, 2/5 animals in the treated groups showed nearly complete abrogation of viral replication and little weight loss by day 7.

### AAV delivery yields therapeutic ACE2 decoy levels in nonhuman primates

Next, we determined which AAV capsid is most efficient at transducing cells of the nonhuman primate (NHP) proximal airways—the desired cellular targets for COVID-19 prophylaxis following nasal delivery of vector. We administered vector using a previously approved intranasal mucosal atomization device (MAD Nasal™), which comprises an atomizing tip with a soft conical nostril seal fit on a standard syringe (Figure 4A). A mixture of 9 AAV serotypes with uniquely barcoded transgenes were administered via the MAD Nasal™ to an NHP. Tissues were harvested 14 days later for evaluation of relative transgene expression using the mRNA bar-coding technique (Figure 4B)(23). The novel Clade A capsid (AAVrh91) we isolated from macaque liver performed best in the nasopharynx and septum (Figure 4C and 4D) with low but detectable expression levels in large airways and distal lung (Figure S6A-G). Clade E and F capsids performed better than AAVrh91 in some non-target tissues such as distal lung (Figure S6A-G). The profile of expression from AAVrh91 illustrates relative distribution of transgene expression with proximal airway structures>intra-pulmonary conducting airway>distal lung (Figure 4E).

**Figure 4.**
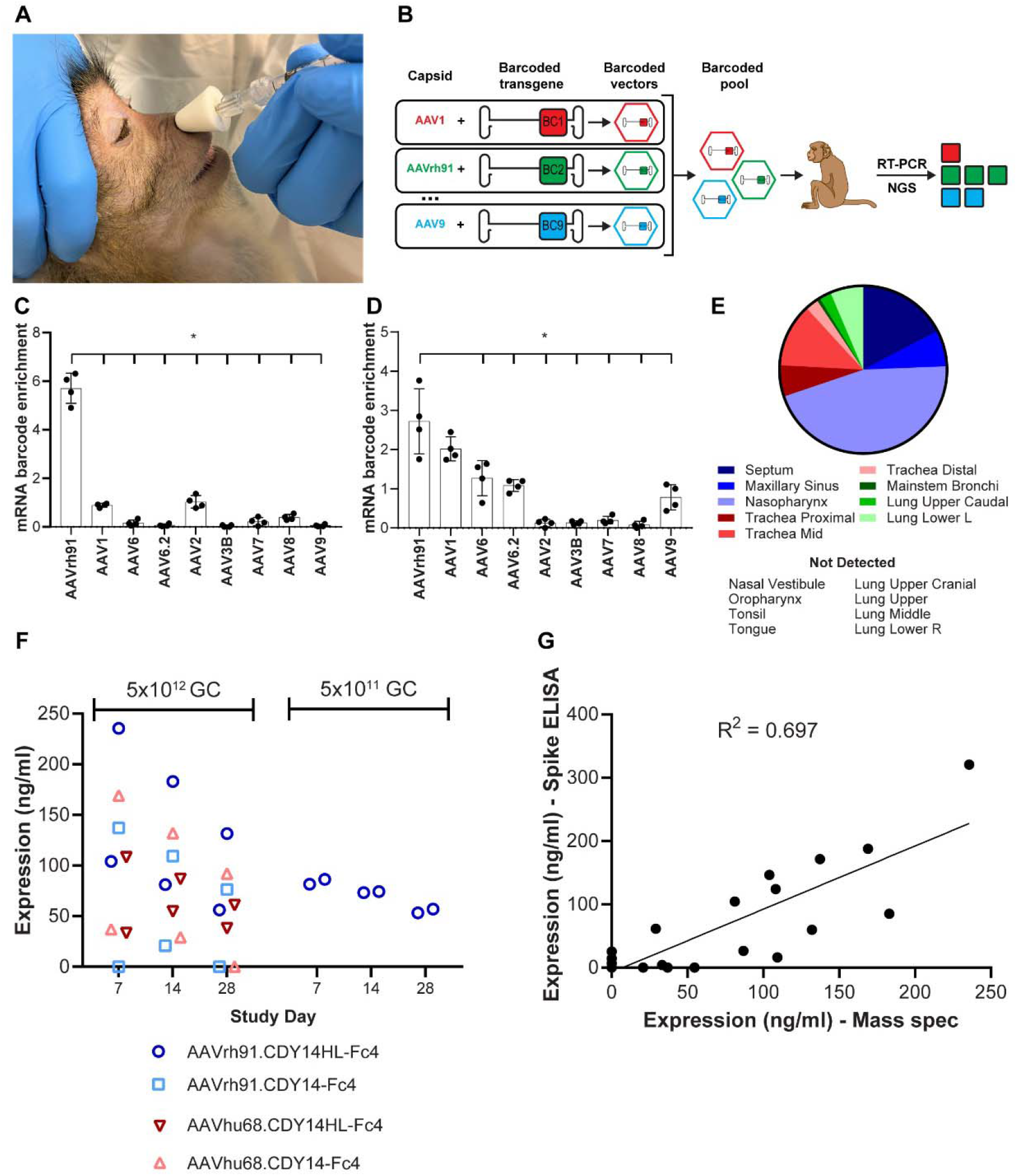
AAV-Delivered Decoy Expression in the Airway of NHP. **A**. NHP vector delivery via MAD Nasal™. **B**. Pooled capsid comparison using mRNA barcoding. mRNA barcode enrichment scores (average across 4 barcodes ± SD) in (**C**) nasopharynx and (**D**) septum. **E**. AAVrh91 transduction profile in airway barcode study. **F**. MS assay of expression in ASF (corrected for BAL dilution) for NHPs (2/group) dosed with ACE2 decoy vector. **G**. Correlation between decoy protein by MS and spike binding by ELISA. Data includes d7 and d14 samples from (F) plus NLF of 3 naïve macaques. We excluded one naïve animal from ELISA analysis because of background binding presumably due to a prior coronavirus infection.

To determine the candidate for clinical evaluation, we conducted a final NHP study where groups of 2 animals were administered 5×10^12^ GC of vectors that differed with respect to capsid (AAVhu68 vs. AAVrh91) and transgene cassettes (CDY14-Fc4 vs CDY14HL-Fc4). NLFs were harvested on days 7, 14, and 28, and animals were necropsied on day 28 for biodistribution. Analysis of pulmonary tissues from day 28 revealed broad distribution throughout the proximal and distal airway, with AAVrh91 demonstrating superior gene transfer to proximal airway structures, as suggested by the barcode study (Figure S6H-I). We estimated decoy protein concentrations in the air-surface liquid (ASF) based on dilution-adjusted MS measurements of NLF (Figure 4F). The effective concentrations at the ASF were in the range that demonstrated neutralization in the *in vitro* assay. Expression was slightly higher with AAVrh91 vs. AAVhu68, and CDY14HL-Fc4 vs. CDY14-Fc4. A subset of samples evaluated for binding to the spike protein of SARS-CoV-2 showed a good correlation with decoy protein as measured by MS. This indicates that the decoy protein produced *in vivo* in proximal airways is indeed functional (Figure 4G).

Based on these data, we selected a candidate for subsequent clinical evaluation called GTP404. This candidate utilizes AAVrh91 as the capsid because of its transduction profile and CDY14HL-Fc4 as the transgene because it retained broad and potent neutralizing activity in the setting of an ACE2-disabling mutation. In preparation for IND-enabling studies, we administered GTP404 at a 10-fold lower dose to two additional NHPs (Figure 4F). Impressive levels of decoy protein were present in nasal ASF with concentrations only slightly reduced in comparison to those achieved with the higher dose.

## Discussion

The rapid emergence of more dangerous and transmissible variants of SARS-CoV-2 in this pandemic is troubling, but not unexpected. The immunological pressures on the virus during natural infections, following antibody therapies, and active vaccines have promoted the emergence of variants(24). It appears that SARS-CoV-2 improved fitness through mutations that both increased affinity for ACE2 and decreased neutralization by antibodies elicited to precursor strains of the virus(15, 25, 26). The density of ACE2 in the nose and airway has been linked to pathogenicity and transmissibility of SARS-CoV-2(27). The lower levels of ACE2 in the proximal airways of children may be responsible for the lower infection rates and milder symptoms in this group(28). It has been proposed that SARS-CoV-2 variants achieve greater infection and transmission through increased affinity(25, 29); here we confirm increased affinity for ACE2 in SARS-CoV-2 strains under positive selection during 2020.

Our original goal in engineering the ACE2 decoy was to improve its potency against SARS-CoV-2, which we accomplished through affinity maturation against the Wuhan-Hu-1spike protein in a yeast display system. Our selection strategy yielded a decoy with high binding and neutralizing activity against a full range of SARS-CoV-2 variants, including B1.1.7 and B1.351, which emerged from the UK and Republic of South Africa, respectively. However, we were surprised to see equally potent binding and neutralization against other betacoronaviruses, including SARS-CoV-1, which was responsible for the 2003 SARS pandemic. The presence of several second-shell mutations in the affinity-matured decoy may contribute to this breadth since the majority of the ACE2 contact surface was preserved (Figure S3). The engineered decoy may be the Achilles’ heel of any ACE2-dependent CoV whose primary driver of fitness – higher binding to its receptor – should further enhance the potency of the ACE2 decoy.

We focused on IN delivery of AAV to express CDY14HL-Fc4 to prevent COVID-19. We used the previously described hACE2-TG mouse challenge model to demonstrate efficacy of the decoy *in vivo*. Treated animals lost less weight, showed reduced lung pathology, and showed less replication of the challenge virus. Our results are consistent with the use of this model to evaluate convalescent plasma(21), protease inhibitors(30), and monoclonal antibodies(31), where weight loss, pulmonary pathology, and viral load were decreased, but not completely abrogated.(21) We believe that the mouse challenge model underestimates the potential efficacy of IN AAV-CDY14HL-Fc4. The dose of SARS-CoV-2 that results in human infection is likely much lower, and therefore, easier to neutralize than the inoculating dose used in the mouse challenge model (2.5×10^6^ particles or 280 PFU). We used a novel AAV Clade A capsid called AAVrh91 to maximize transduction in the proximal airways of NHPs. At a relatively low dose (5×10^11^ GC), we achieved levels of CDY14HL-Fc4 in the ASF that should be sufficient to neutralize SARS-CoV-2 variants. Based on our previous studies—using AAV to deliver broadly neutralizing antibodies against influenza—we believe expression should be durable for at least six months and can be effectively readministered(3-6),(32).

The emergence of three lethal and highly contagious CoV outbreaks in two decades – SARS in 2003, MERS in 2012, and COVID-19 in 2019 – suggests that CoVs will remain a threat to global health. Surveillance of potential zoonotic sources of these CoVs, such as bats, revealed reservoirs of related viruses capable of evolution and cross-species transmission(33). One possible therapeutic application of CDY14HL-Fc4 is in the prevention and treatment of future outbreaks caused by new CoVs that utilize ACE2 as a receptor. GTP404 could be rapidly deployed from stockpiles to contain the initial outbreak and the CDY14HL-Fc4 protein can be leveraged to improve outcomes in those who are infected. The CDY14HL-Fc4 products may be useful in the current COVID-19 pandemic if SARS-CoV-2 variants confound current treatment and prevention strategies. An immediate application could be in immune-suppressed individuals who do not respond to traditional vaccines, develop chronic infection with SARS-CoV-2, and may be reservoirs for new variants(24). The advantage of vector-expressed decoy in preventing COVID-19 infections in immune-suppressed individuals is that this therapy does not rely on the recipient’s adaptive immune system to be effective.

## Materials and Methods

### Yeast display

We generated mutagenized ACE2 gene fragments by error prone PCR using the Diversify PCR Random Mutagenesis Kit (TakaraBio) at multiple mutation levels, mixing the PCR products. We used Gap-repair cloning and high-efficiency LiAc transformation(34) to assemble the ACE2 gene fragments into a centromeric plasmid. The plasmid contained an upstream Aga2 gene fragment, a downstream HA epitope tag with flexible GSG linkers, and was driven by an inducible GAL1 promoter, and contained a low-copy centromeric origin, similar to pTCON2(35). Following transformation or sorting rounds, we passaged the libraries at 10X diversity 3 times in SD-trp before inducing in log phase for 24 hrs at 30°C in SG-CAA(35). For FACS, we stained the yeast with recombinant CoV2 RBD-His6 (Genscript) at diminishing concentrations through the rounds (5nM, 1nM, 0.1nM +/- extended washes of up to 17 hrs for off-rate sorting). We followed up with Mouse anti His6 (Genscript), rabbit anti-HA-PE (Cell Signaling Technology), and goat anti mouse-488 secondary antibody (ThermoFisher) all in phosphate buffered saline (PBS) with 0.1% BSA. Libraries were sorted and analyzed for RBD binding and ACE2 expression at the Penn Flow Core on BD Influx and BD FACSAria II instruments. We extracted plasmid from clones or pools of clones using the Yeast Plasmid MiniPrep Kit (Zymo) and transformed this into bacteria for amplification.

### Next-Generation Sequencing and Analysis

We performed 2×250 paired-end Illumina sequencing on randomly sheared and size-selected ACE2 amplicons from the yeast display rounds. After removing adapters and low-quality reads, we mapped clean reads no shorter than 200bp to the WT ACE2 nucleotide sequences using NovoAlign (v.4.03.01). We translated in-frame sequences with mapping quality scores no lower than 30 and without indels into amino acid sequences and compared to the WT ACE2 protein sequence. We tallied non-synonymous changes for each codon across the ACE2 sequence (18-615). We calculated mutation rates at each codon as follows: (sum of non-synonymous AAs)/(sum of all AAs).

### Expression of ACE2 variant IgG Fc4 fusions and RBDs

We sub-cloned candidate ACE2 synthetic DNA or decoy sequences from the yeast display format into pCDNA3.1, using the endogenous ACE2 signal peptide and appending a human IgG4 Fc domain (residues 99 to 327 from Uniprot reference sequence P01861) and a C-terminal His6 tag. To generate protein for screening we transiently transfected HEK293 cells with plasmid DNA in six-well plates using PEI and collected and clarified supernatant 72 hours later. We quantified expression using the IgG4 Human ELISA kit (Invitrogen BMS2095) with IgG4 standards provided in the kit. For CDY14-Fc4 and CDY14HL-Fc4, we produced the protein in a similar manner but purified it on protein A sepharose followed by dialysis and SDS-Page analysis. We determined the concentration using the predicted extinction coefficient at 280nm. We cloned the synthetic sequences (IDT gBlocks) of RBD [Spike amino acids 330-530 (CoV2), 317-516 (CoV1), and 318-517 (WIV1-CoV)] into pCDNA3.1 between an IL2 signal peptide plus Gly-Ser and a C-terminal His6 tag. We transfected RBD plasmids into HEK293 cells using PEI and collected supernatants 72 hrs later for clarification, concentration, and purification on Ni-NTA resin, followed by dialysis into PBS. We confirmed purity using Coomassie-stained SDS-PAGE analysis. We determined concentrations of the RBD using predicted extinction coefficient at 280nm.

### RBD Binding with SPR

We performed SPR binding analysis using a Biacore T200 instrument (GE Healthcare) at room temperature in HBS-EP(+) buffer (10 mM HEPES pH 7.4, 150 mM NaCl, 3 mM EDTA, and 0.05% P20 surfactant, Cat# BR100669, Cytiva) using a protein A/G derivatized sensor chip (Cat# SCBS PAGHC30M, XanTec Bioanalytics). We injected WT ACE2-hFc4 or CDY14-hFc4 diluted to 60nM in HBS-EP(+) at a flow rate of 10 µL/min for 3 min to capture ∼1,000 response units (RU) on the sensor surface in each cycle. We measured binding of various SARS-CoV RBD proteins to this surface at concentrations ranging from 200 nM to 0.195 nM. RBD binding was measured at a flow rate of 30 µL/min, with a 3 min association time and a 15 min dissociation time. We performed regeneration between binding cycles using 10 mM glycine pH 1.5 injected at a flow rate of 60 µL/min for 1 min. KD values were determined for each interaction using kinetics parameter fitting in the Biacore T200 Evaluation software. We used a global 1:1 binding model and did not adjust for refractive index shift. Data presented are the averate of two or more replicates were measured for each RBD domain tested.

### CoV Pseudotyped Lentiviral Neutralization Assay

We obtained non-replicating lentivirus pseudotyped with CoV spike proteins from Integral Molecular. The reporter virus particles encoded a renilla luciferase reporter gene. We set up neutralization reactions with 100 ul of inhibitor diluted in full serum media and 10 ul of reporter virus. After 1 hour at 37°C, we added 20,000 cells/well in 50 ul of a HEK 293T cell line overexpressing ACE2 (Integral Molecular) and incubated the cells for 48 hours. We measured reporter virus transduction activity on a luminometer (BioTek) using the Renilla Glo Kit (Promega) following manufacturer’s instructions. For higher throughput screens of neutralizing potency, we used crude expression supernatant (described above) in the neutralization assay at 1 or 2 dilutions (typically 10- or 100-fold). We transformed the luciferase reading to an estimated potency (EP) using the following formula: EP = (L*[decoy])/(1-L), where L is the fractional luciferase level as compared to a mock sample (no inhibitor), and [decoy] is the concentration of the decoy in the neutralization well. This was sufficient to rank clones without performing a full titration.

### AAV Vector Production

The University of Pennsylvania Vector Core produced recombinant AAV vectors as previously described (36, 37).

### Decoy Quantification by Mass Spectrometry

#### Standards

Soluble hACE2Fc (produced in-house) was spiked at different levels (0.5-500 ng/mL) into PBS or NLF acquired from a naïve rhesus macaque. Samples were denatured and reduced at 90°C for 10 minutes in the presence of 10mM dithiothreitol (DTT) and 2M Guanadinium-HCl (Gnd-HCl). We cooled the samples to room temperature, then alkylated samples with 30mM iodoacetamide (IAM) at room temperature for 30 minutes in the dark. The alkylation reaction was quenched by adding 1µL DTT. We added 20mM ammonium bicarbonate to the denatured protein solution, pH 7.5-8 at a volume to dilute the final Gnd-HCl concentration to 200mM. Trypsin solution was added at ∼4ng of trypsin per sample ratio and incubated at 37°C overnight. After digestion, formic acid was added to a final of 0.5% to quench digestion reaction.

#### LC–MS/MS

We performed online chromatography with an Acclaim PepMap column (15 cm long, 300-μm inner diameter) and a Thermo UltiMate 3000 RSLC system (Thermo Fisher Scientific) coupled to a Q Exactive HF with a NanoFlex source (Thermo Fisher Scientific). During online analysis, the column temperature was regulated to a temperature of 35°C. Peptides were separated with a gradient of mobile phase A (MilliQ water with 0.1% formic acid) and mobile phase B (acetonitrile with 0.1% formic acid). We ran the gradient from 4% B to 6% B over 15 min, then to 10% B for 25 min (40 minutes total), then to 30% B for 46 min (86 minutes total). Samples were loaded directly to the column. The column size was 75 cm x 15 um I.D. and was packed with 2 micron C18 media (Acclaim PepMap). Due to the loading, lead-in, and washing steps, the total time for an LC-MS/MS run was about 2 hours.

We acquired MS data using a data-dependent top-20 method for the Q Exactive HF; we dynamically chose the most abundant not-yet-sequenced precursor ions from the survey scans (200–2000 m/z). Sequencing was performed via higher energy collisional dissociation fragmentation with a target value of 1e5 ions, determined with predictive automatic gain control. We performed an isolation of precursors with a window of 4 m/z. Survey scans were acquired at a resolution of 120,000 at *m*/*z* 200. Resolution for HCD spectra was set to 30,000 at *m*/*z*200 with a maximum ion injection time of 50 ms and a normalized collision energy of 30. We set the S-lens RF level at 50, which gave optimal transmission of the *m*/*z* region occupied by the peptides from our digest. We excluded precursor ions with single, unassigned, or six and higher charge states from fragmentation selection.

#### Data processing

We used BioPharma Finder 1.0 software (Thermo Fischer Scientific) to analyze all data. For peptide mapping, we used a single-entry protein FASTA database to perform searches. The mass area of the target peptide was plotted against the spike concentration to complete a standard curve.

#### Selection of target peptide

Based on initial *in silico* studies, we selected four peptides as possible sequence-specific matches for targeted quantification. We evaluated sensitivity performance for quantification of the four peptide targets in the NLF background matrix. Following blank injections to establish system cleanliness, replicate injections (n = 3) were made at all levels, from 0.5 ng/mL to 500 ng/mL. Three of the peptides were detected with ANHYEDYGDYWR providing the greatest response across the whole range. We determined retention time (RT) reproducibility across all samples (n = 24) and determined peak area reproducibility and quantification accuracy for each level. Excellent linearity was observed for the levels tested with typical R^2^ > 0.94 for ANHYEDYGDYWR. For ANHYEDYGDYWR, we observed excellent precision and accuracy at all levels, with all replicates within 10% CV. For test articles, 1x or 10x NLF and/or bronchoalveolar lavage fluid (BAL) is treated as previously described without any dilution or protein precipitation. The mass area of target peptide in test articles was compared to the linear calibration generated for the spiked material to determine the level of decoy present in the test article.

### ASF dilutions from serum and lavage urea

We used the urea concentrations in BAL or NLF and in serum collected at the same time to determine the dilution that the lavage introduced to the ASF(38). We quantified urea in mouse BAL and serum, and in NHP NLF using the Urea Assay Kit (Abcam). We obtained serum urea concentrations from NHP from the blood urea nitrogen as part of standard bloodwork lab panels (Antech).

### Spike Binding ELISA

SARS-CoV-2 Spike Protein RBD (Sinobio #40592-V08H) was immobilized on a 96-well plate (0.25ug/mL in PBS, 100ul/well) at 4°C overnight. Plates were then washed 5x with PBS/0.05% Tween and blocked with PBS/1.0% BSA for 1 hour with shaking. Samples (2x dilution in PBS/2.0% BSA) and standards (soluble hACE2-Fc at starting concentration 100ng/ml, 12-point, 1:2 serial dilution, plus a 0.0ng/ml blank, in PBS/1.0% BSA) were added at 100ul/well in duplicate and incubated for 2 hours at room temperature with shaking. Wells were washed as described and biotin conjugated goat anti-human IgG (Jackson AffiniPure #109-065-098; 1:30,000 or Southern Biotech 2049-08; 1:1,000) in PBS/1.0% BSA detection antibody was added to the wells at 100ul/well and incubated for 2 hours at room temperature with shaking. Wells were washed as described, followed by the addition of 100ul/well Streptavidin-HRP (Abcam #ab7403; 1:30,000) in PBS/1.0% BSA for 30 minutes with shaking. Wells were washed as described and incubated in 100ul/well TMB substrate (Seracare #5120-0076) in the dark at room temperature with shaking until reaction was stopped with 100ul/well TMB Stop Solution (Seracare #5150-0021). Absorbances were read at 450 nm using a Spectramax M3 plate reader. We exported and analyzed the data in GraphPad Prism Version 9.0.2. All raw data was blank subtracted. We plotted a standard curve of soluble hACE2-Fc, and the X-axis (concentration) was log_10_ transformed. We performed a 4-parameter nonlinear regression upon the transformed standard curve, and interpolated sample concentrations.

### Determination of Matrix Interference in BAL and NLF Samples

Soluble hACE2Fc was spiked into NLF (0.0, 0.5, 2.0, and 10.0 ng/ml) acquired from a naïve rhesus macaque on the same plate with a standard curve (soluble hACE2Fc starting concentration 100ng/ml, 12-point, 1:2 serial dilution, plus a 0.0ng/ml blank) in PBS/1.0% BSA. We performed the spike binding assay and data analysis as described above.

### Expression study in mice

All animal procedures were performed in accordance with protocols approved by the Institutional Animal Care and Use Committee of the University of Pennsylvania. C57BL/6J mice were purchased from The Jackson Laboratory. Anesthetized mice received an IN administration of 10^11^ GC of AAVhu68.CDY14-Fc4, AAVrh91.CDY14-Fc4, AAVhu68.CDY14HL-Fc4, or AAVrh91.CDY14HL-Fc in a volume of 50 µL or the same volume of vehicle control (PBS) on day 0. On day 7, mice were euthanized and BAL was collected (1 ml of PBS administered intratracheally).

### hACE2 TG Mouse Study

To evaluate the prophylactic efficacy potential of AAV expressing hACE2 receptor decoys against SARS-CoV-2, BIOQUAL, Inc. (Rockville, MD) conducted a challenge study using hACE2 TG mice (Stock No: 034860, The Jackson Laboratory). Mice were administered with either vehicle or 10^11^ GC of AAVhu68.CDY14HL-Fc4 IN on day −7 as described above. On day 0, mice were administered with mock or the SARS-CoV-2 challenge (50 µl of 2.8×10^2^ pfu of SARS-CoV-2, USA_WA1/2020 isolate [NR-52281, BEI Resources]). Mice were euthanized on either day 4 or 7 via cervical dislocation. BAL was collected as described above and aliquoted for viral load assays into Trizol LS (Thermo Fisher Scientific, Waltham, MA) or heat inactivated (60°C for 30 minutes) for decoy protein expression. The lung was collected and split for histopathology into 10% neutral buffered formalin or snap frozen for viral load analysis. RNA extraction for RT-qPCR, the quantitative RT-PCR assay for SARS-CoV-2 RNA, and subgenomic RNA were performed as described(39).

### Histopathology of Collected Organs

The organs collected at necropsy were trimmed and routinely processed for hematoxylin and eosin (H&E) staining. Slides were blindly evaluated by a blinded pathologist using a severity score of 0 (no lesions observed), 1 (minimal), 2 (mild), 3 (moderate), 4 (marked) and 5 (severe) for each finding.

### Intranasal capsid comparison by AAV barcoding

We generated a set of custom barcoded plasmids using degenerate nucleotides that anneal immediately downstream of the stop codon in a GFP reporter construct that contains the AAV2 ITRs. We produced barcoded AAV vectors for each serotype in the study separately by transfecting HEK293 cells as described(36), replacing the typical single ITR-containing plasmid in the transfection mix with an equimolar mixture of 4 uniquely barcoded reporter constructs. We pooled the individual vector preps on an equimolar basis using their digital droplet PCR titers. We determined the absolute barcode distribution in the AAV pool by deep sequencing; we extracted AAV genomes from the pool and performed linear-range PCR using primers that flank the barcode region to generate an amplicon for paired-end Illumina sequencing.

### NHP studies

Rhesus and cynomolgus macaques were obtained from Primgen (PreLabs). NHP studies were conducted at the University of Pennsylvania or Children’s Hospital of Philadelphia within facilities that are United States Department of Agriculture-registered, Association for Assessment and Accreditation of Laboratory Animal Care-accredited, and Public Health Service-assured. For the barcode study, 4×10^12^ GC of the pool AAV preps was delivered IN in a total volume of 0.28 ml to an adult male rhesus macaque using the MAD Nasal™ device. After 14 days, we collected airway tissues at necropsy, and extracted total RNA using Trizol Reagent (Thermo Fisher). We generated cDNAs using Superscript III reverse transcriptase (ThermoFisher) and an oligo dT primer. We used the cDNAs to prepare barcode amplicons for Illumina sequencing as described above. We extracted the relative barcode abundances in input (AAV mixture) and output (tissue cDNAs) from Illumina data. The ratio of output to input relative abundances for each barcode in each tissue is proportional to the relative efficiency of the capsid linked to that barcode in that tissue. Agreement among the 4 barcodes assigned to each capsid allows us to assess assay noise, and detect rare, tissue-specific effects of the barcode itself on transcript stability (none detected). For each tissue, we quantified the total capsid-derived transcript per ug of total RNA using qPCR with a primer/probe set common to all the barcoded reporters.

For the decoy expression in NHPs, cynomolgus macaques (n=2/vector) were administered IN with 5×10^12^ GC of AAVhu68.CDY14, AAVrh91.CDY14, AAVhu68.CDY14HL, or AAVrh91.CDY14HL as described above. An additional two NHPs were administered with 5×10^11^ GC of AAVrh91.CDY14HL. All NHPs were negative for pre-existing neutralizing antibody titres to the administered AAV capsid prior to study initiation (Immunology Core at the Gene Therapy Program). Animals were monitored throughout the in-life phase for complete blood counts, clinical chemistries, and coagulation panels by Antech Diagnostics (Lake Success, NY). On days 7, 14, and 28 NLF was collected (animals placed in ventral recumbency with head tilted to the right, up to 5 mL of PBS delivered in 1mL aliquots, and fluid collected via gravity). Animals were necropsied on day 28 and a full histopathological evaluation was performed.

### Ethics Statement for Study Conducted at BIOQUAL (hACE2 TG Mouse Challenge Study)

This research was conducted under BIOQUAL Institute Institutional Animal Care and Use Committee (IACUC) approved protocol number 21-005, in compliance with the Animal Welfare Act and other federal statutes, and regulations relating to animals and experiments involving animals. BIOQUAL is accredited by the Association for Assessment and Accreditation of Laboratory Animal Care International and adheres to principles stated in the Guide for the Care and Use of Laboratory Animals, National Research Council. Animals were monitored twice daily for clinical signs (specifically ruffled fur, heavy breathing, lethargy) and weighed daily.

### Statistical analysis

Statistical analyses performed using *R* (version 4.0.0). Statistical tests described in figure legends.

## Acknowledgments

We thank Hailey Shankle, Henry Hoff, Shiva Shrestha for notable contributions to the decoy development and characterization. We thank Nathan Denton for assistance with manuscript preparation and graphics. We thank Kirsten Copren and Maggie Shaw for sequencing and analysis support. We thank Victoria Kehm for histology processing and James Tarrant for histopathology analysis. We thank the Immunology Core and the Program for Comparative Medicine of the Gene Therapy Program at the University of Pennsylvania for study support. We thank the Flow Cytometry and Cell Sorting Resource Laboratory at the University of Pennsylvania for cell sorting. All vectors were produced by the Penn Vector Core.

## Funding

This work was funded by the Harrington Discovery Institute, the NHLBI Gene Therapy Resource Program (75N92019D00016), and internal Gene Therapy Program resources.

## Author Contributions

J.J.S. – conceptualization, investigation, methodology, project administration, writing-original draft, writing-review and editing; J.A.G. – formal analysis, methodology, project administration, writing-review and editing; K.T.M. – methodology, resources, writing-original draft, writing-review and editing; S.L. – investigation; R.A.M. – investigation; R.M. – investigation; K.B.T. – investigation, methodology; K.N. – resources; C.D. – methodology, resources; C.H. – conceptualization, methodology; M.H. investigation; H.Y. – formal analysis; X.H. – formal analysis, software; S.J.C. – investigation; J.M.W. – conceptualization, funding acquisition, methodology, supervision, writing-original draft, writing-review and editing.

## Conflict of Interest Statement

J.M.W. is a paid advisor to and holds equity in Scout Bio and Passage Bio; he holds equity in Surmount Bio; he also has sponsored research agreements with Albamunity, Amicus Therapeutics, Biogen, Elaaj Bio, FA212, Janssen, Moderna, Passage Bio, Regeneron, Scout Bio, Surmount Bio, and Ultragenyx, which are licensees of University of Pennsylvania technology. J.M.W., J.J.S, C.H., J.G., M.H., K.N., K.B.T., and S.J.C. are inventors on patents/patents filed by the University of Pennsylvania.

## Data Availability Statement

All datasets presented in this study are included in the article/Supplementary Material.

## List of Supplementary Materials

Figures S1-S6

## Notes

### Summary of Updates

Supplemental files added

