## Supplemental data for "Intranasal gene therapy to prevent infection by SARS-CoV-2 variants"

Figures S1 to S6

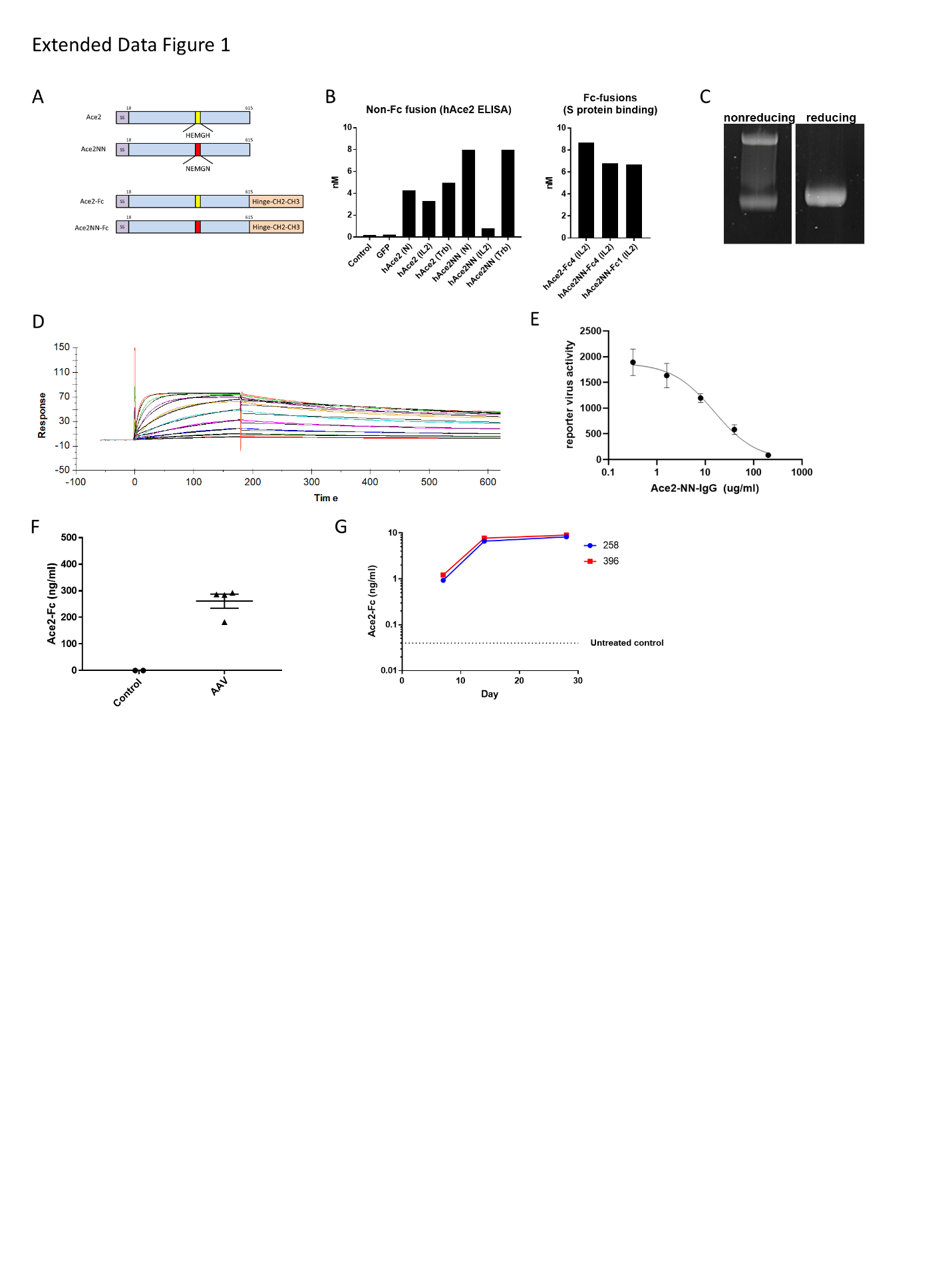
**Supplementary Figures**

**Figure S1. Design and characterization of initial ACE2 decoy construct. A**. Schematic representation of the initial ACE2-Fc4 decoy constructs. ACE2 decoy constructs contained the extracellular domain of ACE2 (amino acids 18-615) with one of three candidate signal peptides (IL-2, native, or thrombin). In some constructs two catalytic histidine residues were mutated to asparagine to abrogate enzymatic activity (designated NN). Constructs were designed with no Fc, or the Fc domain of IgG1 or IgG4. **B**. Constructs were expressed in HEK293 cells and detected in supernatant using a sandwich ELISA to ACE2 (for constructs without an Fc domain) or an ELISA with SARS-CoV-2 spike protein as a capture antigen and an anti-human IgG polyclonal antibody for detection (for Fc fusion proteins). **C**. The candidate ACE2-NN-Fc4 fusion protein was expressed in HEK293 cells, purified by protein A chromatography, and analyzed by SDS PAGE under reducing and nonreducing conditions **D**. The affinity of the purified ACE2-NN-Fc4 decoy protein for monomeric spike protein in solution was quantified by Biacore SPR. k_on_ = 2.6 x 10^5^ M^-1^ s^-1^, k_off_ = 0.00093 s^-1^, t_1/2_ = 745 s, K_D_ = 3.5 nM , R_max_ = 67 RU. **E**. The purified ACE2-NN-Fc4 protein was titrated against Wuhan CoV2 pseudotyped lentivirus bearing a luciferase reporter. The IC50 was obtained from a fit of these data (15 ug/ml) **F**. The candidate construct (ACE2-NN-Fc4) was packaged in an AAV vector (hu68 capsid) and administered IN to WT mice. Seven days after administration, BAL was collected for measurement of transgene expression using an ELISA with SARS-CoV-2 spike protein as a capture antigen to confirm that the decoy receptor expressed *in vivo* was functional. BAL from similar experiments was 6-fold diluted from the ASF as determined by comparison of BAL and serum urea. Thus, we determined that ASF concentrations of the decoy were likely below 2 ug/ml. **G**. Two NHPs (IDs 258 and 396) received 9 x 10^12^GC of an AAVhu68 vector expressing a soluble ACE2-NN-Fc fusion protein via the MAD (Figure 4). Nasal lavage samples were collected weekly after vector administration and concentrated 10-fold for analysis. The concentration of the decoy receptor in NLF was measured by MS. Urea measurements in similar experiments indicate that 10X nasal lavage is ~8-fold diluted from ASF. We therefore determined that ASF concentrations of the decoy were less than 100 ng/ml.

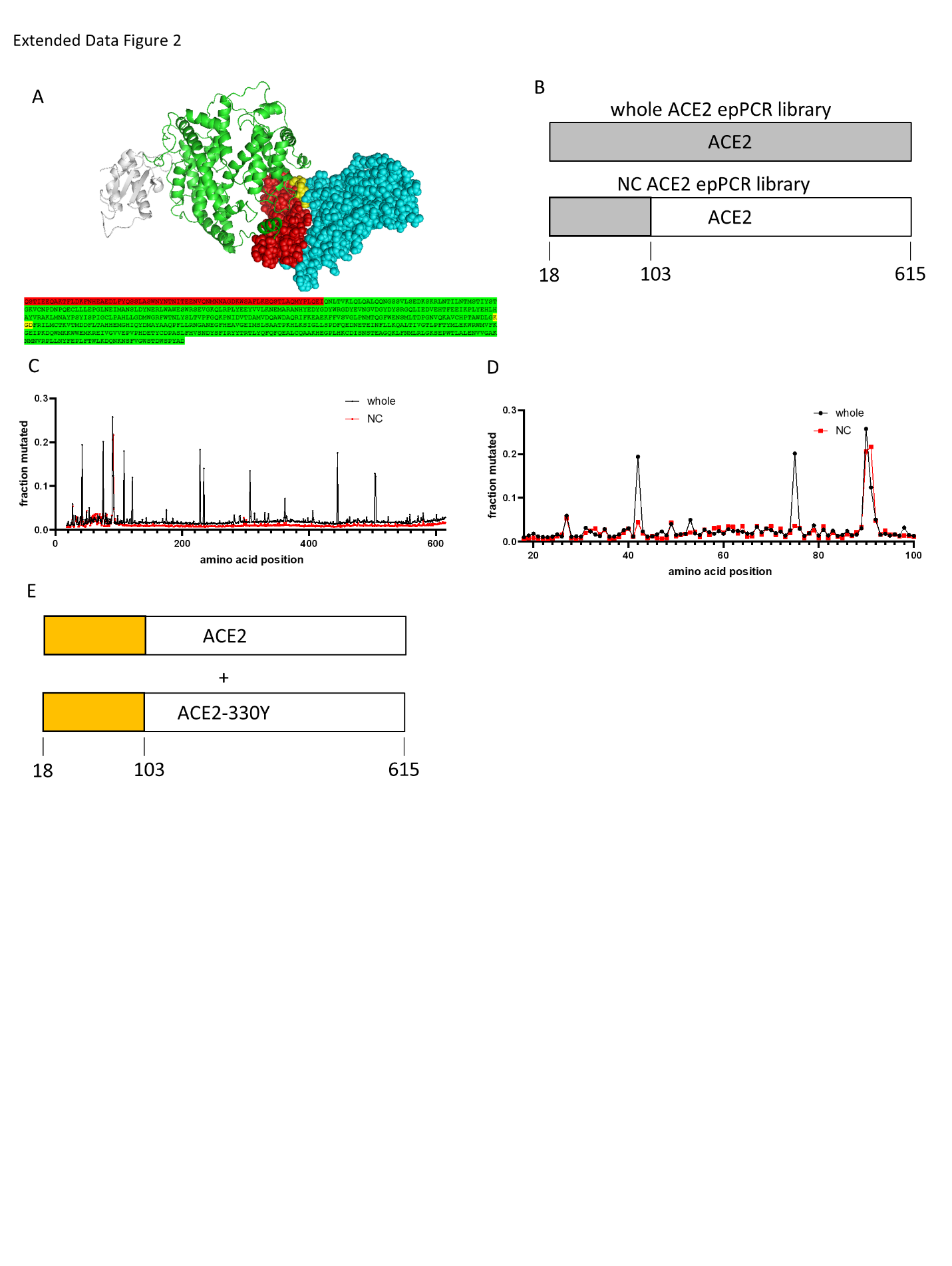

**Figure S2.** Design and selection of primary and secondary yeast display libraries. **A**. Structure of CoV-2 RBD (blue spheres) bound to human ACE2 (green ribbons, red and yellow spheres) (6M17.pdb (*41*)). Most ACE2 contacts with RBD are limited to the amino acids 18-88 (red spheres) and a patch of amino acids that are more C terminal (yellow spheres). The gene sequence for ACE2 is shown below in the same coloring. **B**. We designed two primary yeast display libraries: 1) the whole ACE2 gene fragment was mutagenized (Whole) and 2) the mutagenesis was limited to only the first 96 amino acids (NC) to concentrate the mutagenesis on the region most likely to impact RBD binding. The regions shaded gray were subjected to error-prone PCR to introduce mutations. **C**. Deep sequencing of yeast display plasmids extracted from the final round of sorting for the Whole and NC libraries. The fractional rate of mutation at each position in 18-615 of ACE2 is plotted. Improving mutations occurred mostly in the first 96 amino acids regardless of the input library. These include RBD contact residues, second-shell residues, and the distal consensus N-glycan site at position 90, an apparent negative regulator of RBD binding. Though C-terminal mutation load was generally associated with poor ACE2 expression (data not shown), several consensus C-terminal mutations emerged from the whole ACE2 primary sorts. These include a substitution to Y at position 330, which we identified in a clone with improved binding. **D**. A detailed plot of the mutational frequencies in Whole and NC library final round sorts for residues 18-100. The libraries yielded many of the same mutants in this region with improved binding activity. **E**. Schematic representation of secondary library design. We isolated 300 yeast colonies from the sorted primary (Whole and NC) libraries, analyzed them individually for RBD binding and ACE2 expression by flow cytometry, and selected 90 isolates with validated binding improvements. Next we generated a secondary library by shuffling selected ACE2 genes using the staggered extension process (StEP) method(*9*). Given that most improving mutations were N-terminal, we shuffled only residues 18-103 of the input templates (orange shaded region in the schematic), matching these with a mixture of unmutated and N330Y C-terminal DNAs in a multi-fragment assembly yeast transformation.

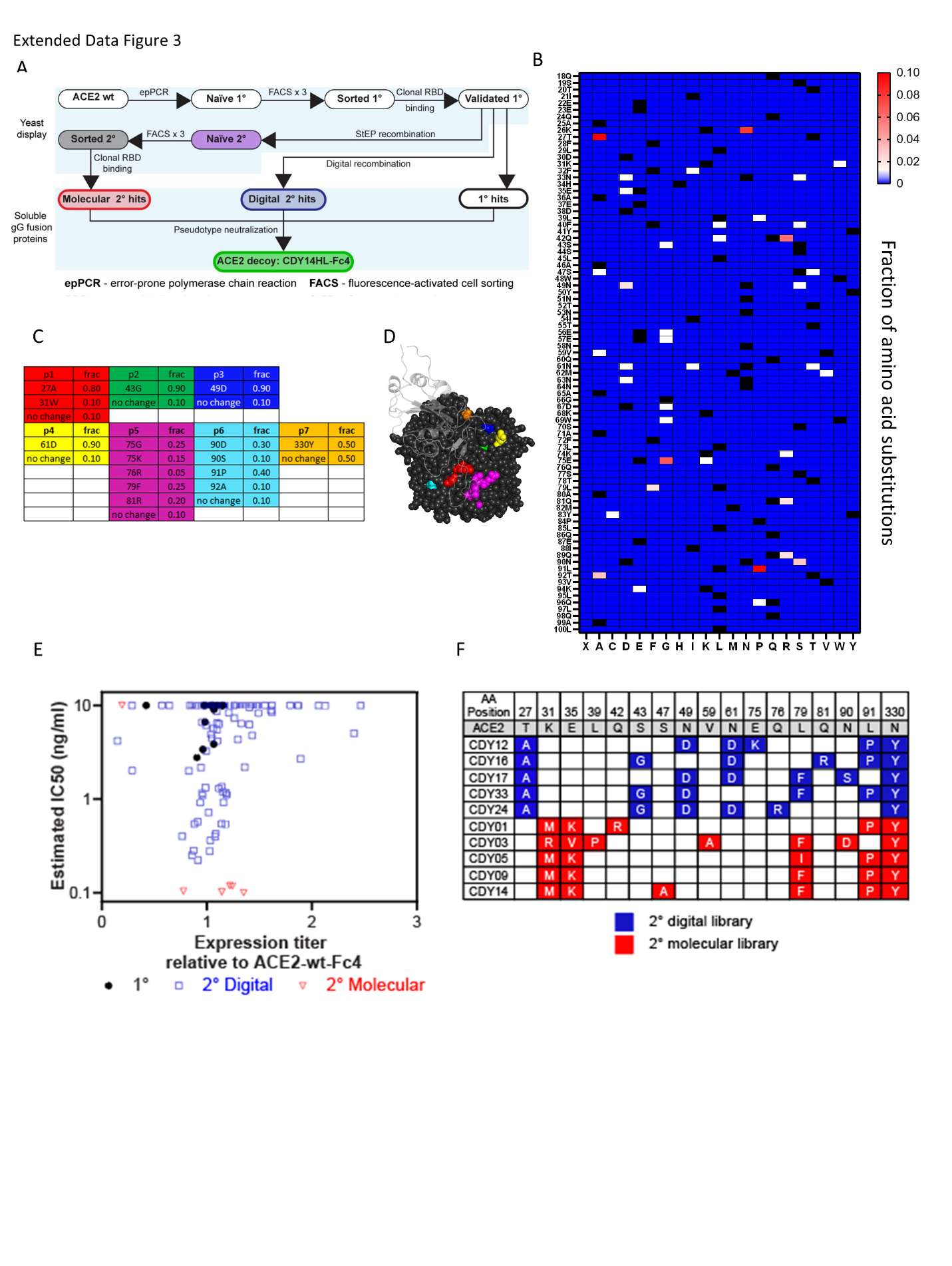

**Figure S3. A.** Schematic of parallel paths to the generation of affinity matured ACE2 decoy. After generating improved RBD binding sequences from a primary round of sorting, we undertook a parallel path to digitally recombine the most frequent mutations in addition to continued diversification and sorting. Primary library hits, digital recombinants of those hits and isolated clones from the second, more stringent round of yeast display sorting were all cloned as Fc4-fusion proteins and screened in a CoV2 pseudotype neutralization assay. **B**. We selected 300 clones from primary yeast display library sorts for clonal RBD binding analysis using flow cytometry in the yeast display format and selected 90 clones with validated binding improvements. A pool of plasmid DNA from those 90 isolates was subjected to deep sequencing for mutational analysis, and the rates of all possible amino acid substitutions are presented in this heat map by amino acid position. Black squares represent the wt amino acid at each position. **C**. The goal was to generate a collection of synthetic (digital) recombinants of the observed mutations in this data set. We grouped subsets of the validated mutations into 7 regions (p1 -p7) and assigned frequencies to the mutations based loosely on observed frequencies in the data set, allowing for the wt residue at 10% or 50% depending on the position. We biased the mutation selection towards second-shell positions to avoid directly remodeling the ACE2:RBD contact positions where possible. **D**. In order to maximize the chance of mutations working together productively, we chose the groupings in (C) based on 3D structure (6M17.pdb (*40*)) such that direct contacts between groups would be minimized. We randomly selected primary screen mutation combinations based on the frequencies in (C), and had these digital recombinants synthesized for cloning as secreted IgG Fc4 fusion proteins. **E.** We screened primary yeast display hits, digital recombinants, and secondary yeast display hits in a pseudotyped lentivirus reporter assay for CoV-2 neutralization at one or two dilutions from the expression supernatant, noting the expression titer relative to ACE2-wt-Fc4 control. **F**. Mutations associated with the 5 best digital recombinant hits and the 5 best hits from a secondary round of yeast display sorting.

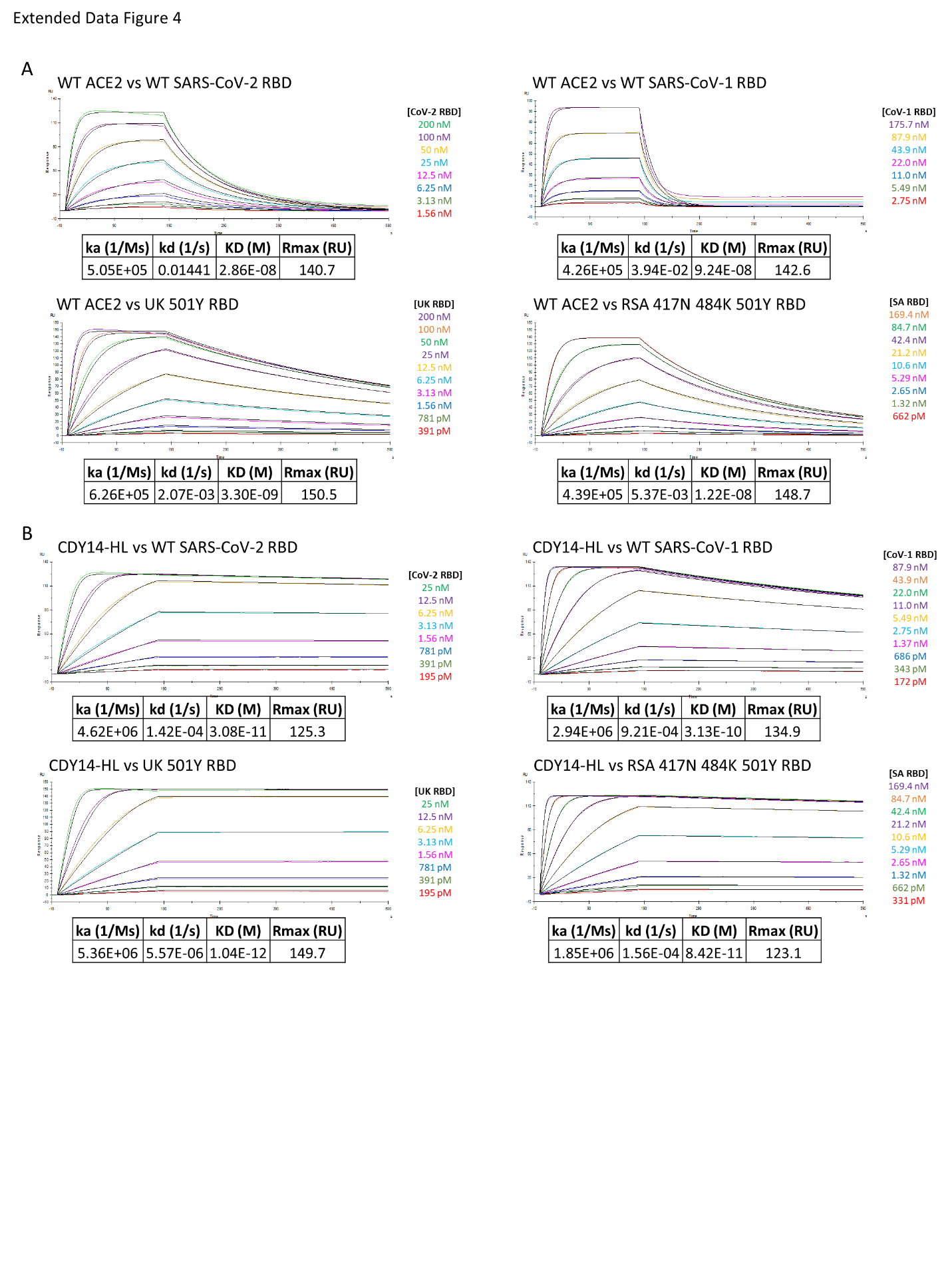

**Figure S4.** Example SPR data for RBD binding assay. **A**. Raw data (colored lines) and fits (black lines) for immobilized ACE2-wt-Fc4 on the SPR chip surface binding injected RBDs (concentrations listed). Parameters of the fits, including the dissociation equilibrium constant (KD) are listed below each panel. These data contributed to Figure 2B and 2C. **B**. Raw data (colored lines) and fits (black lines) for immobilized CDY14HL-Fc44 on the SPR chip surface binding injected RBDs (concentrations listed).

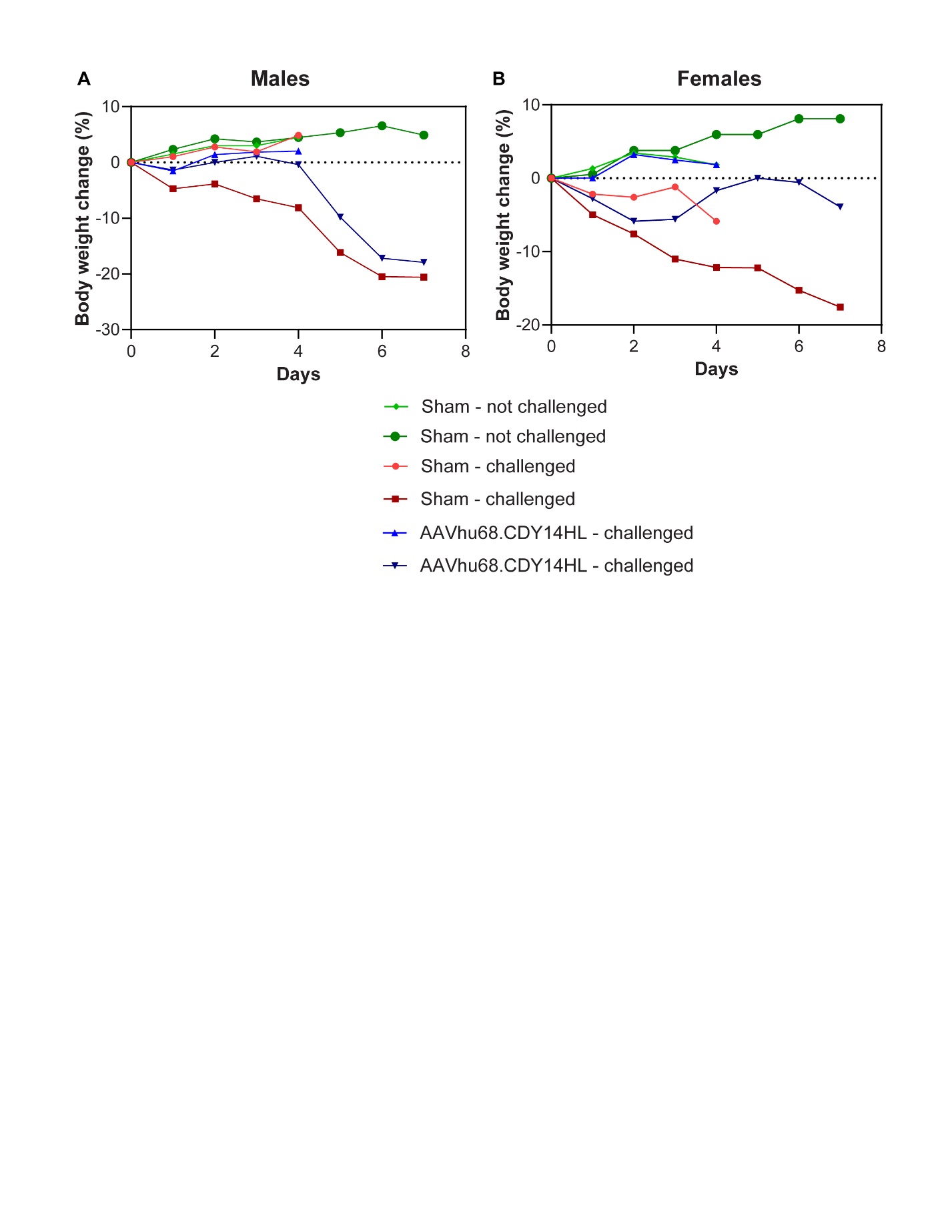

**Figure S5.** Challenge study in mice. Average weight loss (percentage) in males **A**. and females **B**. hACE2-TG mice that received Challenge Placebo and Challenge Decoy.

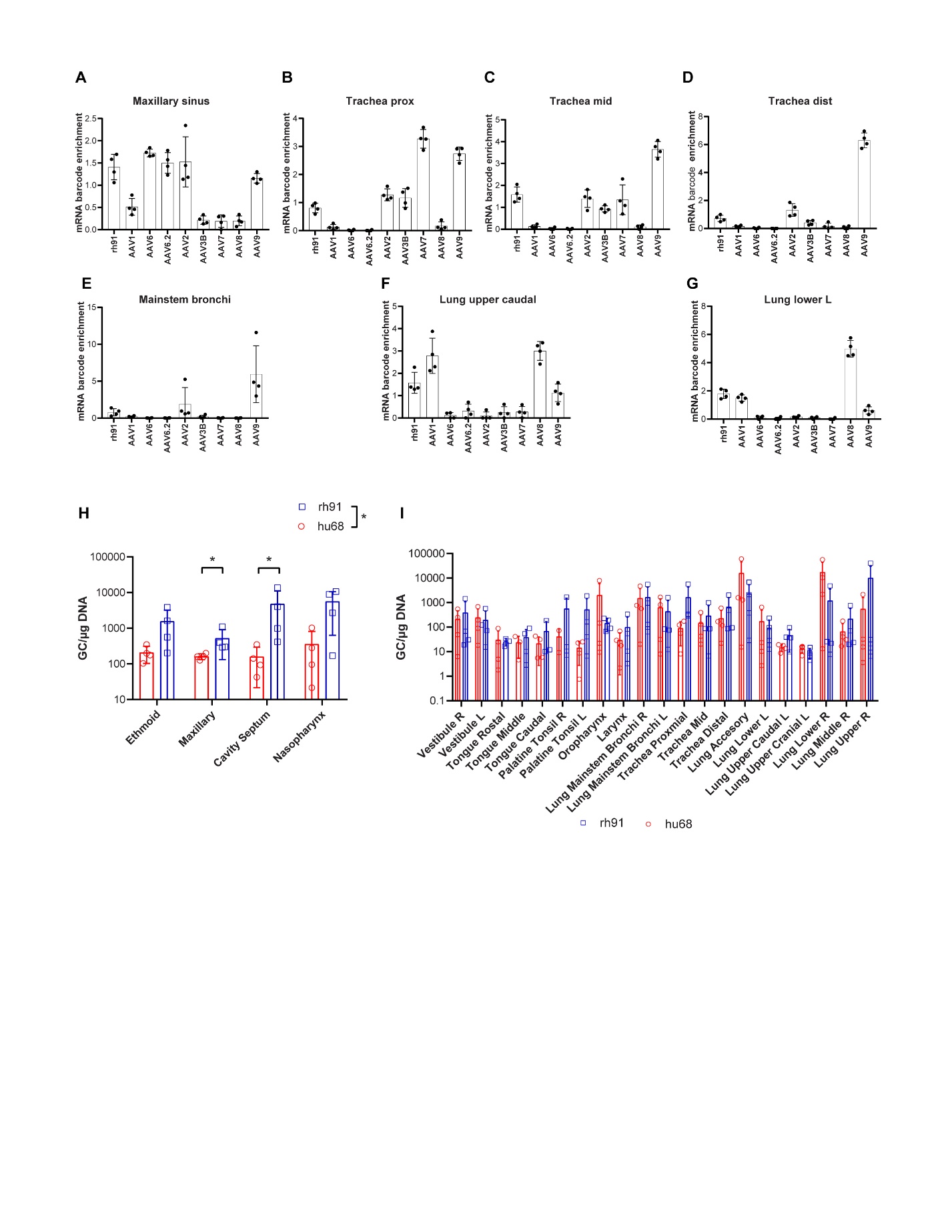

**Figure S6.** AAV capsid selection for NHP IN delivery. **A**-**F**. mRNA barcode enrichment in airway tissues for a mixture of 9 barcoded serotypes delivered IN at 2.7E11 GC each. We assigned four uniquely barcoded transgenes to each capsid at manufacture. Data show the enrichment score (tissue abundance in RT-PCR-NGS/ injection mixture abundance in PCR-NGS) for all 4 barcodes per capsid with mean and SD. **H and I**. Four NHP were IN dosed with rh91 or hu68 vectors encoding decoy transgenes at 5E12 GC. Data show the biodistribution of vector genomes in airway tissues 28 days after dosing. Rh91 achieved higher gene transfer in upper airway tissues (H), particularly in the maxillary sinuses and cavity septum. Gene transfer in lower airway tissues was more variable (I).
